# Genetic distance between complex repeats

**DOI:** 10.1101/443721

**Authors:** Luca Ferretti, Aurora Ruiz-Herrera, Alice Ledda

## Abstract

Complex nucleotide or aminoacid repeats with long units play an important role in proteins. The evolutionary analysis of these variants is challenging due to genetic diversity within repeat units as well as variability in the arrangement of different units along the repeat sequence. Here we present a new approach for the computation of genetic distances between complex repeats. This method takes into account evolutionary processes including point mutations, insertions and deletions of repeat units, as well as duplication of single units. We provide an algorithm for the computation of these distances along with the corresponding global pairwise alignment of repeats. As an example, we apply our approach to the evolution of repeat units in the highly polymorphic zinc-finger repeat domain of the PRDM9 protein across wild populations of house mice. This approach opens the way for new insights into the evolutionary history of polymorphic repeats.

## Introduction

Reconstructing phylogenies of repetitive regions has always been a challenge for evolutionary biologists. In the dawn of the genomic era these regions were masked in reconstructing phylogenies due to the intrinsic difficulties they posed [4]. The evolution of repetitive regions is mostly shaped by peculiar genomic processes like non-homologous recombination, replication slippage etc [5]. These processes often play a major role in the evolution of such regions compared to point mutations. This increases the difficulties in modelling the evolution of such genomic regions, as these processes need to be included in realistic approaches [1].

Existing models are focused on specific types of repeats such as microsatellites, Copy Number Variants etc, where repeat units are similar or identical and the only degree of freedom is their length in terms of the number of repeated units. A more challenging and richer family of repeats is represented by complex satellite repeats with long repeat units (tens of bp or aminoacids) with some degree of internal variability between similar units, as well as variability in the repeat composition in terms of groups of similar units. Only a few of the existing bioinformatic tools are able to deal effectively with this kind of repeats, and most of them provide alignment of protein repeats [9, 10, 8]. To our knowledge, the only practical approach with a currently available implementation is repeat-aware Multiple Sequence Alignment of such repeats with ProGraphMSA+TR [13].

On long-term scales, all units of a repeat often derive from a single ancestral unit via point mutations and duplication/slippage. This is true even for complex repeats with different types of units [1]. However, on short evolutionary scales (between close species or within species), insertions and deletions of highly diverged units represent clearly different processes with respect to recent duplication/slippage followed by divergence via point mutations. Existing methods for repeat alignment do not discriminate between these processes, and the difference between these processes is not taken into account in the computation of the genetic distance.

Here we develop a new approach for the computation of genetic distances for complex repeats. Our approach includes point mutations, insertions and deletions of whole repeat units, as well as duplications of single repeat units as a consequence of non-homologous recombination, slippage or related biological processes. Single-unit duplications and indels can have different weights and therefore contribute differently to the genetic distance. We also present a version of the Needleman-Wunsch algorithm [7] to compute the genetic distance between pairs of repeats, as well as their optimal pairwise global alignment according to the new genetic distance defined here. The result of this algorithm is a modified alignment that allows for further evolutionary analyses on single-unit duplication/slippage.

We illustrate the potential of our approach by studying the alignments and genetic distances between sequences of the zinc-finger repeat domain of the PRDM9 protein from multiple populations of *Mus musculus*[14]. These sequences are highly polymorphic and they appear to be under complex selection.

## Methods

In our approach, repeat units are treated as fundamental blocks. Only processes preserving the integrity of each block are considered. The identification of repeat units is outside the scope of this paper. The choice of the repeat units is left to the user.

We assume that all repeat units share a high sequence similarity. Hence, Multiple Sequence Alignment of all units from all repeats can be easily obtained from any alignment method. It is therefore straightforward to compute pairwise genetic distances *μ*(*u, u*′) between units *u* and *u*′. Many possible definitions of genetic distance between biological sequences exist; the choice of the most appropriate one is left to the user.

### Definition of distance

We define the genetic distance between repeats as an edit distance, i.e. as the minimum cost to change a repeat to another through a series of elementary operations. The cost is defined as the sum of the weights of all elementary steps. In our case, elementary operations are inspired by biological processes. They are:

- point mutations, small indels and other within-unit processes (weight *w*_*m*_);
- insertions and deletions of whole units (weight *w*_*i*_);
- single-unit duplication/slippage (weight *w*_*s*_).

The edit distance is minimised by one or possibly several alignments between repeats *r* = (*r*_1_, *r*_2_, *r*_3_ …) and 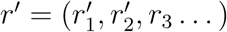. For each of these alignments, we denote by 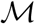 the set of all matching units between *r* and *r*′ (the uth unit in the repeat *r* corresponds to the *m(u*)th unit in *r*′). 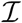 and 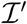 denote the sets of inserted units in *r* and *r*′ respectively. Finally, *Ѕ* denotes the set of duplicated units in *r*, where the *u*th unit is the result of a duplication of the *s*(*u*)th unit (which could be either the (*u*+ 1)th or the (*u*- 1)th unit). *Ѕ*′ and *s*′(*u*) denote the same quantities for *r*′.

Our definition of the distance between *r* and *r*′ is

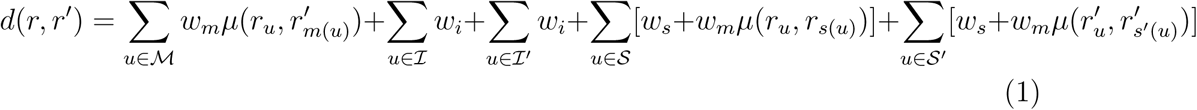

This distance can either be derived from multiple repeat alignment [13] or it can be directly computed using the algorithm discussed in the next section.

### Needleman-Wunsch algorithm with single-unit duplications

We present a modification of the classical Needleman-Wunsch algorithm [7] to include single-unit duplications in the computation of the distance. For each pair of repeats, this algorithm provides their genetic distance as well as the corresponding global alignment.

Our algorithm mirrors closely the standard Needleman-Wunsch algorithm. The differences with respect to the standard version of the algorithm lie in the choice of weights and in the way insertions/deletions are handled.

Our selection of weights is as follows:

- mismatch between units *r*_*u*_ and 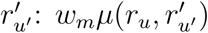
- insertion of a whole unit *r*_*u*_ in repeat *r*:

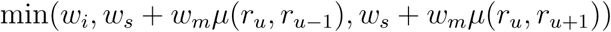

Given these weights, the distance is then computed as in the standard algorithm.

Furthermore, when the best partial alignment results from an insertion in either of the two repeats, we consider the minimum among the values of *w*_*i*_, *w*_*s*_ + *w*_*m*_ μ(*r*_*u*_, *r*_*u*-1_) and *w*_*s*_ + *w*_*m*_ μ(*r*_*u*_,*r*_*u*+1_). These values correspond to three different scenarios:

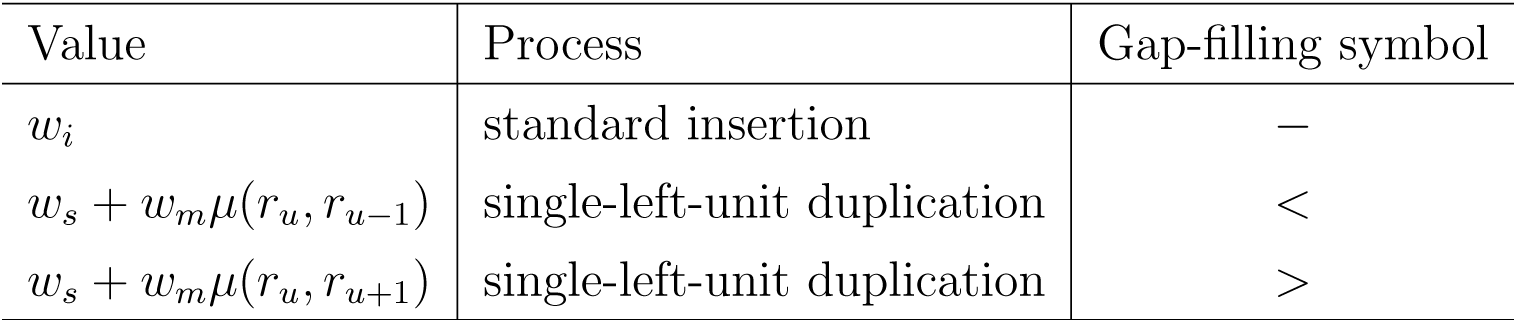

The alignment is computed as in the standard algorithm, but for each insertion, the corresponding gap-filling symbol from the above table is then used to represent the gap in the alignment. In this way, it is possible to separate single-unit duplications from actual insertions/deletions. Note that there could be ties among the above values; in this case, multiple optimal alignments are possible.

## Results

### Single-unit duplications versus insertions/deletions in PRDM9 sequences

To show both the rationale and the impact of single-unit duplications on repeats, we consider a dataset of 208 sequences of the zinc-finger repeat domain of the PRDM9 protein [6]. These sequences are derived from different populations of house mice Mus musculus sampled from Madeira [2, 14]. These populations appear to be highly polymorphic in the PRDM9 locus, with repeats of different length (8 to 16 units) and variable sequence. Each repeat unit is 28 aminoacids in length; three of these aminoacid positions are exceptionally variable among units, probably because of complex selection pressures [3]. We analyse the nucleotide sequences of these repeats both including and excluding the 9 bases corresponding to the hyper-variable loci.

Figure 1 illustrates the role of single-unit duplications. Neighbour repeat units, which are affected by single-unit duplication, tend to have more similar nucleotide sequences than random units within the same repeat or within different repeats (significant by Mann-Whitney U-test, *p* < 10^−15^). This is true both considering all loci or non-hyper-variable ones only. Considering all loci, adjacent repeat units have an average distance of 2.95, versus average distances of 3.44 for random units within the same repeat and 3.35 between units in different repeats. Excluding hyper-variable codons, the average distance drops to 0.79 between neighbour units, compared to 1.11 between units in the same repeat and 1.06 between random pairs of units. The difference between units in the same repeat or in different repeats is small and it is significant only when testing all loci. These results provides support and rationale for modelling single-unit duplication/slippage as an independent process involving adjacent repeat units.

**Figure 1:**
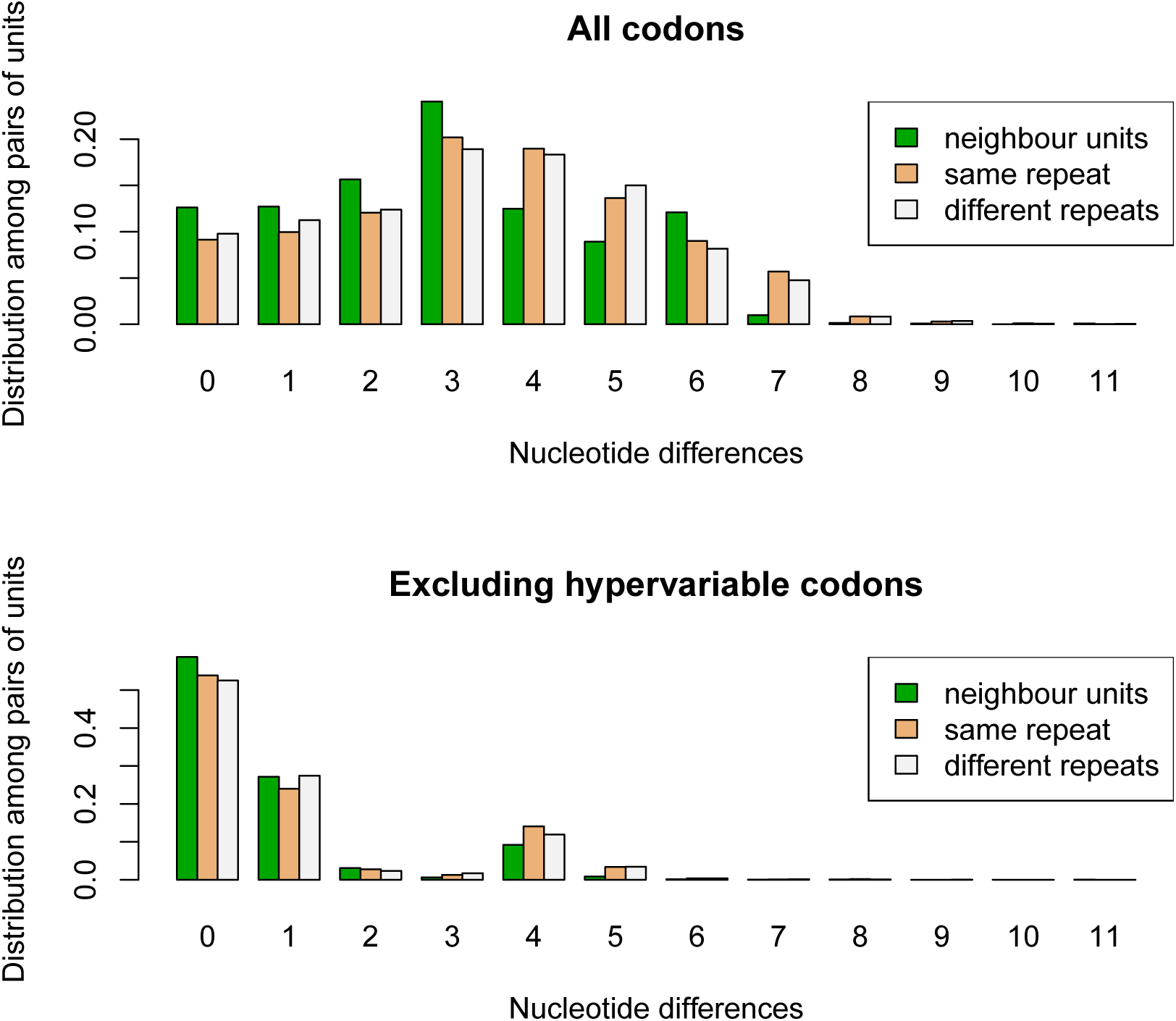
Barplots of the distributions of pairwise nucleotide distances between repeat units. The three distributions are obtained from pairs of adjacent units in the same repeats, any pair of units in the same repeats, and pairs of units in different repeats, respectively.

The effect of modelling single-unit duplication/slippage in the pairwise alignment and in the computation of the distance is shown in Figure 2. The genetic distance proposed in this paper is compared to the simple edit distance derived from the Needleman-Wunsch algorithm with the same parameters but *w*_*s*_ = 0. The distance including single-unit duplication is the lower of the two by definition, but the standard Needleman-Wunsch distance is significantly higher (7.3 versus 5.4 for the choice of parameters favoured in [14] and 5.2 versus 3.6 for the other). This emphasises again the frequent occurrence of single-unit duplication/slippage in this system of polymorphic repeat variants.

**Figure 2:**
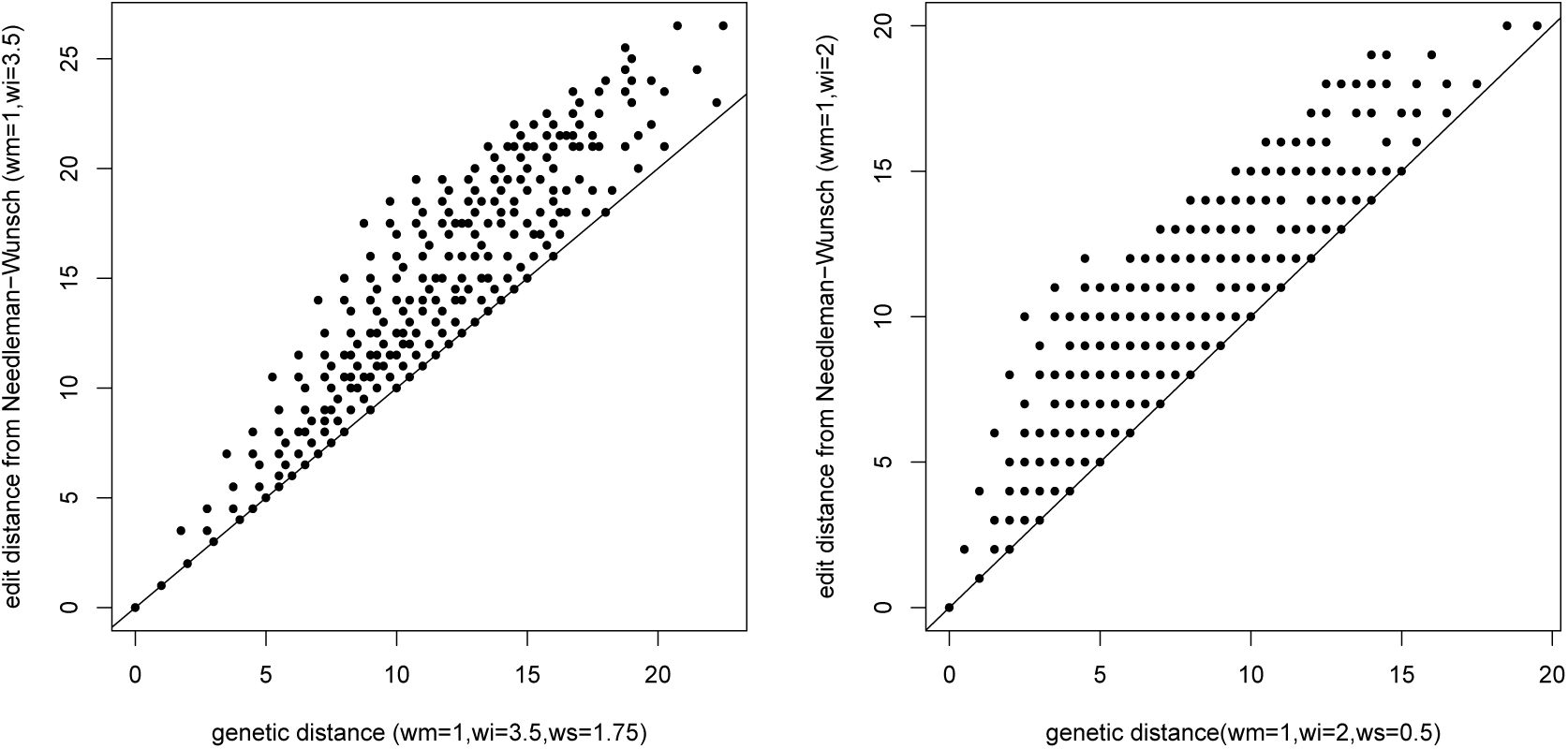
Comparison between genetic distances from our approach and from the classical Needleman-Wunsch algorithm with the same parameters (and *w*_*s*_ = 0). Genetic distances are computed for two different sets of parameters (left: *w*_*i*_ = 3.5, *w*_*s*_ = 1.75; right: *w*_*i*_ = 2, *w*_*s*_ = 0.5), always excluding hyper-variable codons. The parameters on the left correspond to the actual ones chosen in [14] by maximising the agreement between the repeat-based genetic distance and the genetic distance obtained from SNP genotyping data for the genomic region surrounding PRDM9.

In the absence of a gold standard, measuring the improvement of our approach with *w*_*s*_ > 0 is not a trivial task. A reasonable way is to make use of the nature of the biological data. Under the assumption of constant mutation rates, the genetic distances should be proportional to the cophenetic distances along the genealogical tree of the sequences. If the tree is inferred e.g. by Neighbour Joining based on the genetic distance [12], the scale of the tree is the same as the genetic distances, hence a measure of goodness-of-fit is the sum of the squared residuals between genetic and cophenetic distance, normalised by the squared average cophenetic distance. According to this measure, there is not a large different in the goodness of fit for the choice of parameters in the right panel of Figure 2 (2.9 · 10^−2^ versus 3 · 10^−2^), but there is a significant improvement for the more realistic parameters in the left panel (2.8 · 10^−2^ versus 4.3 · 10^−2^). This further supports the inclusion of single-unit duplication/slippage in the modelling and in the genetic distance.

## Discussion

In this paper we present a simple approach to the computation of genetic distances for complex repeats. Phylogenetic reconstruction of their evolutionary history can then be extracted using Neighbour-Joining or other distance-based methods [12]. A byproduct of this approach is an Needleman-Wunsch algorithm for global alignment of repeats. This algorithm can be easily extended to a version of the Smith-Waterman algorithm [11] to provide local alignment of highly divergent repeats.

The key feature of our approach is that we consider single-unit duplications in addition to insertions/deletions. This implies that the method depends on two parameters (*w*_*i*_/*w*_*m*_ for indels and *w*_*s*_/*w*_*m*_ for duplications). More complicate processes could be included, such as duplication from non-adjacent units or large insertions/deletions, at a price of extra parameters to be tuned. A solution could be to add extra features such as separate “gap opening” and “gap extension” penalties instead of a single insertion cost, and similar parameters for duplications. This could improve the handling of large non-homologous recombination events.

A structural limitation of our approach is that complex rearrangements would be captured as alignments with many insertions/deletions, implying large genetic distances. Another limitations is that by treating units as irreducible blocks, we neglect processes involving only subparts of units or shifted units. Processes that affect only a part of the unit sequence, such as within-unit recombination, shifted insertion/deletion/duplication/slippage and so on, are therefore approximated as occurring at one of the extremes of the unit or as involving multiple units. Despite these drawbacks, our method provides a fast and effective estimate of genetic distances for polymorphic repeats from related species or populations.

## Acknowledgments

We thank Philipp Schiffer, Covadonga Vara and two anonymous referees of Molecular Ecology for comments and suggestions. A R implementation of the algorithms described here is provided at https://github.com/lucaferretti/RepeatDistance.

